# Stable Isotope Probing-nanoFTIR for Quantitation of Cellular Metabolism and Observation of Growth-dependent Spectral Features

**DOI:** 10.1101/2024.01.24.576656

**Authors:** David J. Burr, Janina Drauschke, Katerina Kanevche, Steffen Kümmel, Hryhoriy Stryhanyuk, Joachim Heberle, Amedea Perfumo, Andreas Elsaesser

**Author notes:** These authors contributed equally (David J. Burr, Janina Drauschke, Andreas Elsaesser).

## Abstract

This study utilizes nanoscale Fourier transform infrared spectroscopy (nanoFTIR) to perform stable isotope probing (SIP) on individual bacteria cells cultured in the presence of ^13^C-labelled glucose. SIP-nanoFTIR simultaneously quantifies single-cell metabolism through infrared spectroscopy and acquires cellular morphological information via atomic force microscopy. The redshift of the amide I peak corresponds to the isotopic enrichment of newly synthesized proteins. These observations of single-cell translational activity are comparable to those of conventional methods, examining bulk cell numbers. Observing cells cultured under conditions of limited carbon, SIP-nanoFTIR is used to identify environmentally-induced changes in metabolic heterogeneity and cellular morphology. Individuals outcompeting their neighboring cells will likely play a disproportionately large role in shaping population dynamics during adverse conditions or environmental fluctuations. Additionally, SIP-nanoFTIR enables the spectroscopic differentiation of specific cellular growth phases. During cellular replication, subcellular isotope distribution becomes more homogenous, which is reflected in the spectroscopic features dependent on the extent of ^13^C-^13^C mode coupling or to specific isotopic symmetries within protein secondary structures. As SIP-nanoFTIR captures single-cell metabolism, environmentally-induced cellular processes and subcellular isotope localization, this technique offers widespread applications across a variety of disciplines including microbial ecology, biophysics, biopharmaceuticals, medicinal science and cancer research.

## 1. Introduction

Single-cell analyses have made tremendous progress in recent years, providing new insights into fundamental aspects of cell behavior such as differential growth, responses to external stimuli, cell sensing, intercellular communication and interactions. Intercellular heterogeneity (both phenotypic and metabolic) is a consistent feature of cellular populations, ranging from different individuals within clonal populations,^[1]^ multi-species ensembles,^[2, 3]^ and of course multicellular organisms. Even when genetically identical cells are cultured under the same conditions, they may display different phenotypes.^[4, 5]^ Traditional population-based analyses examine a bulk number of cells, relying on the assumption that a clonal pool of microorganisms is homogenous, and as a result, such methods fail to capture variation between single microbial cells that may arise from mutations, spontaneous environmental variation or a combination of the two. Such outlier organisms have the potential to disproportionately contribute to the makeup of a developing population, which serves as an evolutionary strategy, playing a functional role in many biological processes, from cellular differentiation to bet-hedging as an environmental adaptation.^[4, 6]^ Understanding intercellular heterogeneity provides insights into various microbial eco-physiological mechanisms and is of the utmost relevance to explaining dynamics of cell factories, antibiotic resistance, response to drugs, aging and cancer development. To address questions of cell-to-cell heterogeneity and overcome the limitations of traditional population-based analyses, numerous new single-cell technologies have been developed to target the underlying mechanisms of cellular heterogeneity. These cutting-edge techniques include single-cell multi-omics,^[7, 8]^ quantitative imaging,^[9]^ mass spectrometry^[10]^ and vibrational spectroscopy.^[11, 12]^

Infrared (IR) spectro-microscopy is a powerful tool for single-cell chemical studies. The lateral resolution of this technique is however defined by both the long wavelength of IR radiation and the diffraction limit, and thus is unsuitable for resolving subcellular features. In contrast, IR nanoscopy overcomes diffraction-based spatial resolution limits by focusing IR light onto the tip of an atomic force microscope (AFM) and recording the interaction of scattered radiation with matter in the optical near-field, where the object-to-tip distance is less than the wavelength used. The optical resolution is therefore defined by the shape and diameter of the AFM tip, and as such, molecular information can be acquired at remarkable resolutions, down to 20 nm^[13, 14]^ (well below subcellular limits), with an approximate penetration depth of 100 nm.^[15-18]^ By employing a broadband IR laser in an interferometric scheme, IR absorption spectra can be obtained through nanoscale Fourier-transformed infrared spectroscopy (nanoFTIR). Alternatively, scattering-type scanning near-field optical microscopy (sSNOM) scans the sample surface at a single IR frequency, simultaneously acquiring AFM topography and IR-based chemical information, yielding molecularly specific sample imaging. Both sSNOM and nanoFTIR are exceptionally powerful tools for biologically relevant investigations and have been employed in label-free imaging and spectroscopy of systems ranging from well-defined biomolecules^[14, 19, 20]^ to whole cells.^[21]^ Single-cell IR nanoscopy studies have demonstrated oxidative stress of red blood cells,^[22]^ differentiated plant cell wall components,^[23]^ performed spectral observations of live bacterial cells suspended in liquid media,^[24]^ and acquired high-resolution tomography of the internal membrane-bound organelles of microalgae.^[25]^ These recent advancements highlight the versatility and applicability of near-field imaging techniques in life sciences.

Variation in atomic mass intrinsically affects molecular vibrations, thus it is possible to induce observable vibrational spectroscopic shifts by altering the isotopic ratio of composite atoms of biomolecules and subcellular components. Consequently, the cellular uptake of growth media labelled with stable isotopes can serve as an indicator of ongoing metabolic processes. For instance, metabolization of ^13^C provides valuable information on cellular protein and lipid synthesis. Although the majority of cellular stable isotope probing (SIP) studies have been conducted on a bulk number of cells, recent investigations have coupled SIP with techniques capable of single-cell measurements.^[26]^ SIP coupled with nanoscale secondary ion mass spectrometry (nanoSIMS) has provided insights into bacterial metabolic activity,^[27]^ phenotypic heterogeneity,^[28, 29]^ nutrient cycling,^[30, 31]^, and microbial interactions and ecosystem functioning.^[32, 33]^ Metabolic profiling of microorganisms has been achieved at the single-cell level using both SIP-Raman spectroscopy^[34-36]^ and mid-IR photothermal-SIP-fluorescence *in situ* hybridization.^[1]^

To further expand the range of single-cell spectroscopic techniques, here we introduce SIP-nanoFTIR as a novel technique that combines the super-resolution and molecular specificity of nanoFTIR with the integration of detectable stable isotopes into metabolically active cells. High spectroscopic contrast between sample and substrate allows for resolution at the single-cell level. This innovative approach simultaneously acquires nm-resolution AFM imaging in parallel to spectra, providing visualization of cellular topography, phenotypic features and intracellular protein localization. The nanoscale resolution and molecular specificity of this technique can be maximized to great potential by performing subcellular hyperspectral nanoFTIR measurements, allowing research to be conducted on biomolecule composition and identification of metabolic pathways with unprecedented precision. The integration of these complementary methods offers a powerful, comprehensive and versatile toolset for investigating cellular dynamics at the single-cell level, making it an ideal technique for exploring complex biological systems and uncovering the intricacies of single-cell behavior in health and disease. When compared with several of the alternate single-cell techniques described above, SIP-nanoFTIR provides several significant advantages;^[26]^ as SIP-nanoFTIR utilizes a low-energy IR-light source, radiation damage is minimized and cellular samples of interest remain intact throughout the analysis. The non-destructive nature of this technique preserves the integrity of delicate biological structures and enables researchers to perform longitudinal studies without perturbing the cells. Isotopic labelling simply alters compounds that are natively integrated by microorganisms, unlike techniques that rely on foreign chemical markers to identify cellular regions or processes (e.g., fluorescence-based techniques). This leaves the biochemistry unaltered and, thus, cellular replication and metabolic activity are unimpeded.^[37]^

In this study, we performed stable-isotope labelling by supplying the model organism *Escherichia coli* K12 with ^13^C-glucose supplemented growth media. As bacteria readily consume glucose as an energy source, heavy carbon atoms are integrated into newly synthesized amino acids and subsequent biomolecules. Heavy isotope incorporation causes a redshift in IR absorption,^[34, 38]^ and as such, this can be utilized as a means of measuring single-cell metabolic activity. Here, quantification of single-cell protein synthesis was performed based on the observed redshift of the amide I band.^[34, 38]^ Nanometer-scale visualization of biosynthetic activity allowed us to trace the emergence of cell-to-cell metabolic heterogeneity. As cellular translational activity and the physiological state of the cell can be directly influenced by environmental conditions, nutrient limitation resulted in an increase in intercellular heterogeneity and induced cellular morphological changes. In addition, unique changes to the amide region of single-cell spectra were correlated with the specific cellular growth phase of the population. Considering that heterogeneity in protein expression significantly contributes to cell-to-cell phenotypic variability,^[5]^ SIP-nanoFTIR is well-suited for analyses of population dynamics and eco-physiological mechanisms in both unicellular and multicellular systems. As a result, we envision significant interest and diverse applications across various research areas, including microbiology (e.g., the functioning of environmental and host-associated microbiomes, examination of slow growing or non-cultivable yet viable microbes, nutrient cycling, and antibiotic resistance), biotechnology (e.g., bioproduction in cell factories) and medicine (e.g., aging, cancer and cell disease-associated research, and drug delivery systems). The capability of SIP-nanoFTIR to simultaneously localize, visualize and quantify translational activity greatly expands the single-cell analytical toolkit and provides avenues for valuable advancements in understanding the molecular mechanisms underlying cellular behavior and intracellular nutrient localization.

## 2. Results

### 2.1 Principle and function of SIP-nanoFTIR

NanoFTIR is performed by scanning the sample surface with an AFM tip (**Figure 1**), providing the sample topography with nm resolution. Simultaneously, a broadband IR laser is focused on the tip apex, detecting optical information from amplitude and phase of the scattered IR light. The tip oscillates at a frequency (Ω) that allows for signal demodulation at higher harmonics (nΩ), minimizing background contamination of the detected signal. Thereby, far-field scattering intensities are effectively suppressed and signals from the optical near-field are exclusively recorded.

**Figure 1:**
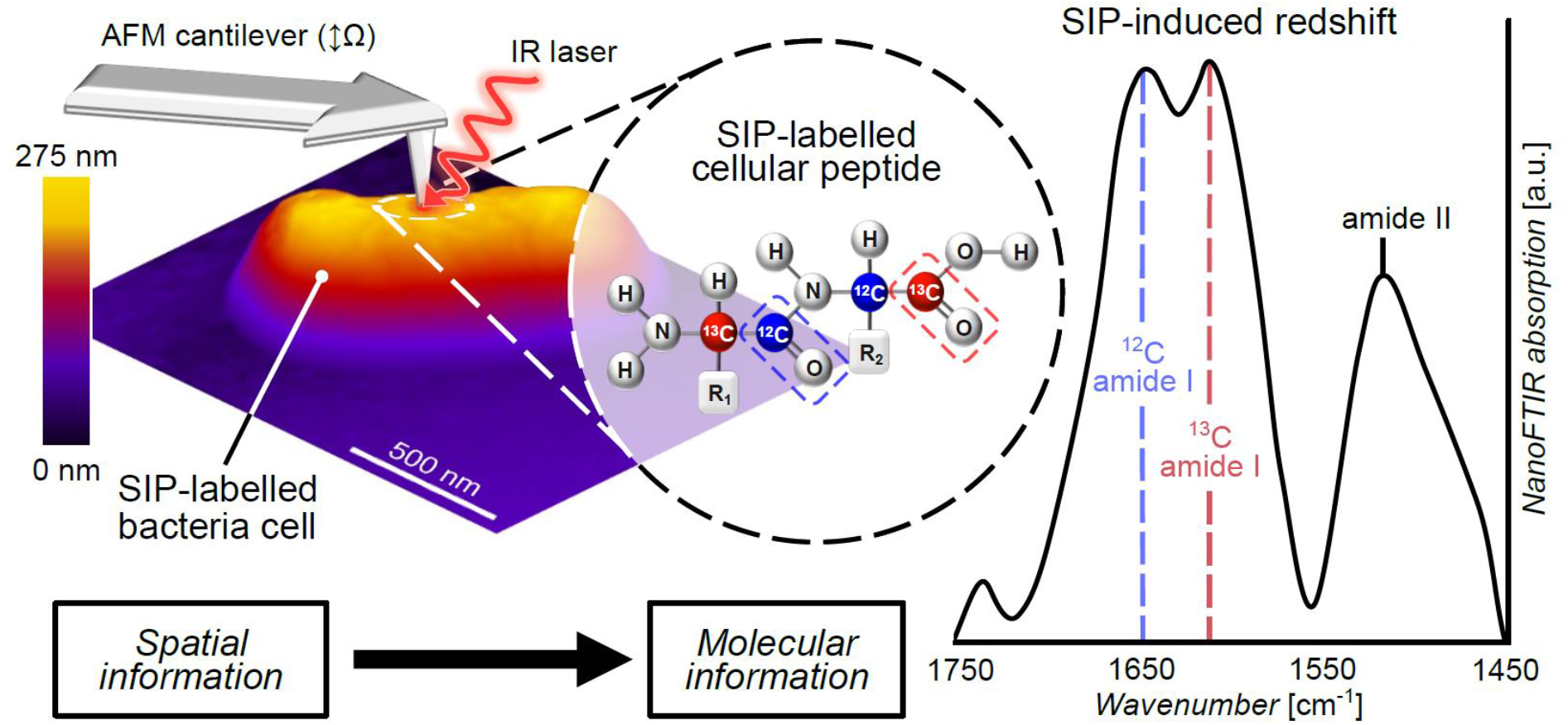
Principle of operation of the SIP-nanoFTIR technique. An illustrative diagram of a single *Escherichia coli* cell cultured under a 50:50 ratio of ^12^C and ^13^C-labelled glucose. Spatial and topographical information is acquired via AFM imaging. IR radiation is focused on the oscillating AFM tip, and scattered light from the sample is detected, providing molecular vibrational information. A cellular peptide molecule is specifically exemplified, being comprised of a 50:50 mixture of ^12^C and ^13^C atoms. An increase in atomic weight reduces vibrational energy, causing an ∼40 cm^-1^ redshift in IR absorption, which results in the distinct dual amide I peaks observed.

^13^C-induced spectral redshifts are only present in IR-absorbance peaks dominated by carbon related vibrations; most prominently, the amide I absorption band, which is primarily due to the C=O stretching vibrations of the peptide backbone. When ^13^C is integrated into cellular proteins, the increase in atomic mass decreases the vibrational energy, and hence the amide I band downshifts by ∼30-40 cm^-1^. In contrast, the amide II peak is primarily derived from the coupling of the C=O stretching with the N-H bending vibrations and as a result, the only potential SIP-induced vibrational-energy changes are due to contributions from neighboring carbon atoms. Therefore, ^13^C inclusion results in only an ∼10 cm^-1^ redshift of the amide II peak that is far more ambiguous to interpret than that of the amide I peak (Figure 1).

A series of *E. coli* cellular standards were cultured to late-exponential phase under specifically defined proportions of ^13^C-labelled glucose. Each of these cellular standards were measured using elemental analyzer isotope-ratio mass spectrometry (EA-IRMS), providing the bulk averaged ^13^C atomic percentage content of each cellular standard (**Table S1**). Although the use of (non-isotopically labelled) chemical fixatives resulted in the reintroduction of a non-negligible portion of ^12^C, this is a well-documented phenomenon.^[27, 39, 40]^ Implementation of this IRMS quality control step provided a means to calibrate the IR-acquired amide I redshift. Therefore, when the non-linear relationship between the proportion of supplied ^13^C-labelled glucose and the true cellular atomic percentage (Table S1) was accounted for, the normalized ^12^C:^13^C peak intensity ratio could be used to quantify the single-cell ^13^C content, with a mean dynamic accuracy of ∼3% (**Figure S1**).

### 2.2 Quantification of single-cell metabolism and subcellular carbon localization

In cultures supplied with increasing fractions of ^13^C-labelled glucose, the amide I absorption peak maximum was observed shifting from 1655 cm^-1^ (±1 cm^-1^) to 1613 cm^-1^ (±2 cm^-1^). Rather than a continuous spectral redshift, a stepwise increase in supplied ^13^C-labelled glucose treatments resulted in a proportional variation of the relative intensities of the ^12^C- and ^13^C-amide I peaks. For instance, cultivation with 50% ^13^C growth media resulted in a distinct and characteristic dual amide I peak shape, each with similar intensities (**Figure 2a**). A ^13^C-induced change was also observed in the amide II peak (1535 cm^-1^), which broadened significantly with increasing ^13^C fraction (Figure 2a). Independent of the proportion of ^13^C provided, an additional shoulder was present ∼20 cm^-1^ lower than the amide I peak. This spectral feature is most prominent at 1630 cm^-1^ and 1590 cm^-1^ in the 0% and 100% ^13^C treatments, respectively, and is less evident in mixed isotope treatments as it is obscured by the double peak structure of the partially red-shifted amide I (Figure 2a). This small shoulder can be attributed to membrane proteins in a β-barrel conformation,^[41]^ which show spectral enhancement in comparison to far-field spectroscopic techniques, due to the <100 nm penetration depth of nanoFTIR spectroscopy.^[15-18]^

**Figure 2:**
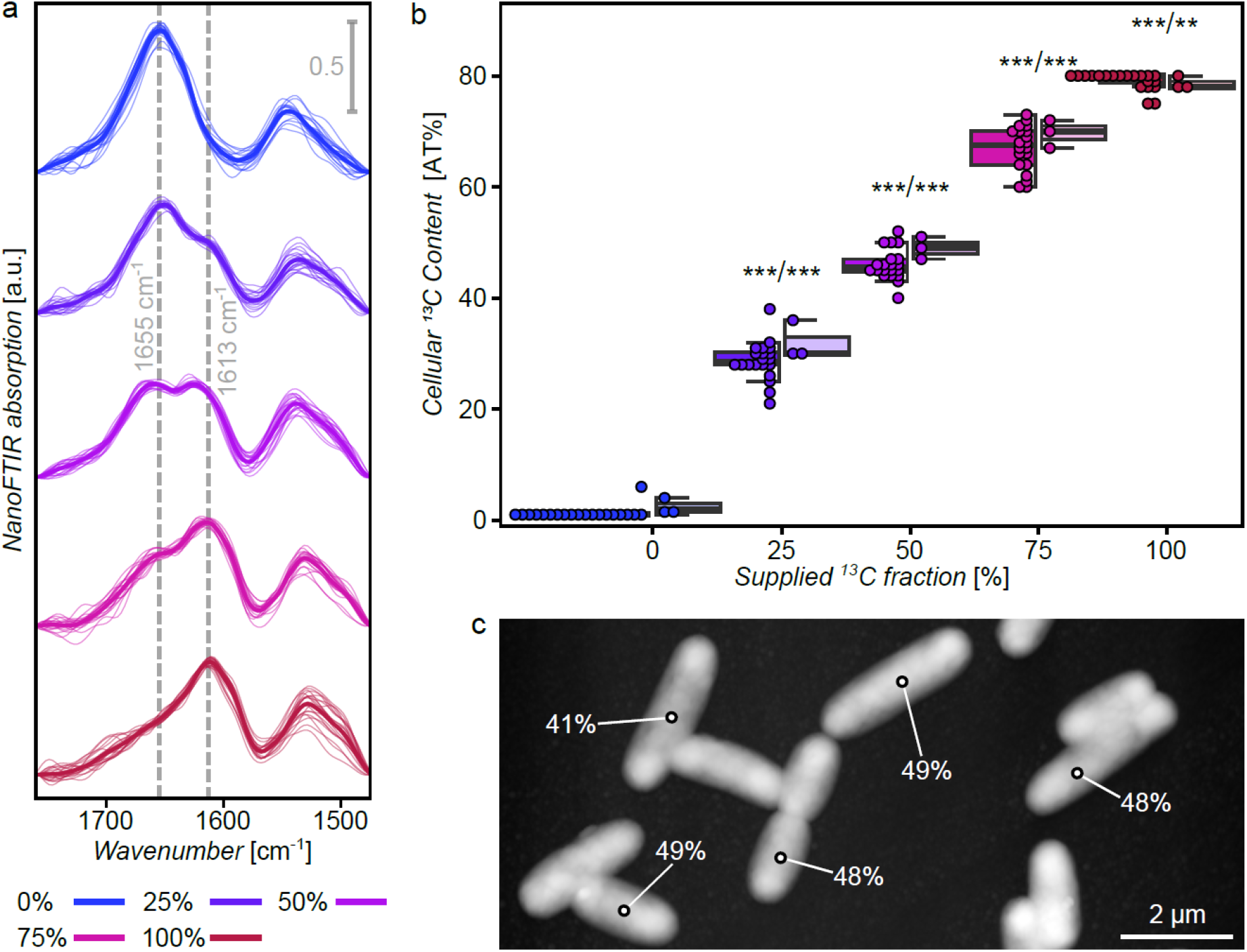
Quantification of single-cell carbon uptake using the SIP-induced redshift. 25% incremental ^13^C-labelled glucose treatments are represented with a blue (0%) to red (100%) color gradient. **a** NanoFTIR absorption spectra (2^nd^ harmonic) of individual single cells, with the mean spectrum of each treatment (n=20, technical replicates) shown in bold. Dashed vertical lines mark the ^12^C-(1655 cm^-1^) and ^13^C-amide I bands (1613 cm^-1^). **b** Back-to-back boxplots (displaying median, upper and lower quartiles, and the maximum and minimum excluding outliers) of quantified single-cell ^13^C content. Dots represent individual data points. NanoFTIR recordings are shown on the left (darker) and IR-reflectance microscopy measurements (lighter) on the right. Asterisks’ indicate statistical differences between treatments (p-value < 0.001 = ***, < 0.01 = **), as determined by a one-way ANOVA at the 95% confidence level. **c** A representative AFM image of the 50% ^13^C-labelled glucose treatment. The cellular ^13^C-content, as detected by nanoFTIR is overlaid, demonstrating the variance in quantified ^13^C content of neighboring single cells. Maximum z-distance = 250 nm.

When the proportional amide I redshift was used to quantify the ^13^C content of individual single cells from five different ^13^C-labelled glucose treatments (n=20, increasing in 25% increments), the mean cellular ^13^C content of each treatment was 1.3%, 28.7%, 46.0%, 67.0% and 79.1%, respectively, with each differing at the 99.9% significance level (**Figure 2b, Table S2**). Comparing single-cell SIP-nanoFTIR measurements were against bulk cell number IR reflectance microscopic measurements, a similar ^13^C-labelled glucose-concentration dependent ∼40 cm^-1^ redshift was observed (**Figure S2**). Upon quantification of IR reflectance microscopy spectra, the mean cellular ^13^C content of each respective treatment (2.3%, 32.0%, 49.0%, 69.7% and 78.7%, respectively) all statistically differed at least at the 95% confidence level (**Table S3**, Figure 2b).

Although the average output of these two techniques were highly similar, quantification at the single-cell level using SIP-nanoFTIR revealed large-scale differences arising within treatments. With a shift from bulk cell number measurements to single-cell observations, a ∼2.5 times increase in data spread is observed across all treatments, with the most prominent differences arising in mixed isotope treatments. Notably, the 50% ^13^C-labelled glucose treatment resulted in observations of single-cell ^13^C content ranging from 40-52% (Figure 2b). Cell-to-cell variance in ^13^C fraction was not spatially correlated, rather, large deviations occurred between neighboring cells. This is demonstrated in **Figure 2c**, where an 8% range of cellular ^13^C content was observed within a single 16x8 μm field of view. Single-cell observations provided a means of verification that changes in the provided fraction of ^13^C were not inducing any morphological differences. Additionally, no isotopically-induced variations in cellular growth occurred (doubling time: F_(1, 2)_ = 0.062, p = 0.826; carrying capacity: F_(1, 2)_ = 3.400, p = 0.207; **Figure S3**).

To examine potential subcellular variations in carbon localization, hyperspectral nanoFTIR mapping was performed, measuring a total of 312 spectra over the surface of a single cell cultured under a 50% fraction of ^13^C. Subcellular spectral differences were observed, with the highest spectral quality being measured in the central regions of the cell (**Figure 3a**), corresponding to the areas of the cell surface closest to horizontal alignment (**Figure 3b**). While both the unreferenced scattered amplitude (**Figure 3c**) and the amide II absorption were relatively homogenous across the entire surface of the cell, the amide I intensity decreased rapidly towards the cell edges (**Figure 3d, e**). This observation can be related to the molecular orientation of C=O oscillators in membrane proteins and the electric field direction of IR radiation scattered by the tip in nanoFTIR. This hyperspectral analysis emphasized that selectively measuring nanoFTIR spectra in the center of each cell is a feasible method to acquire isotopic information representative of the entire cell.

**Figure 3:**
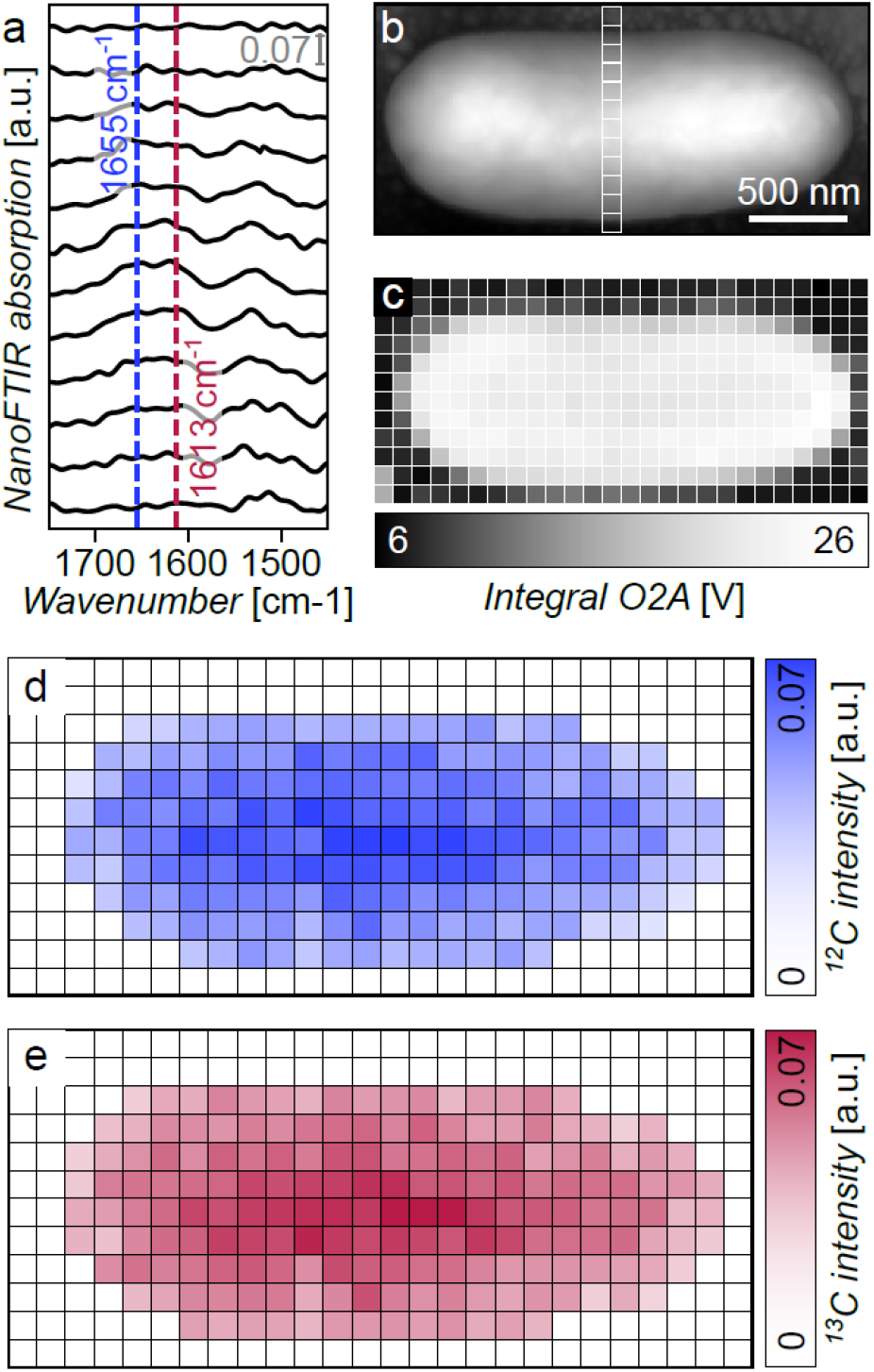
Subcellular hyperspectral SIP-nanoFTIR mapping. **a** 12 exemplary nanoFTIR absorption spectra (2^nd^ harmonic) acquired from positions represented by the transect in Figure 3b. Vertical line represent the ^12^C and ^13^C amide I peak positions, respectively. **b** AFM scan of a 50% ^13^C-labelled *E. coli* cell. Maximum z-distance = 212 nm. **c** Sample masking generated via integrated laser power of unreferenced scattered amplitude. Pixel size = 100x100 nm. **d** 1655 cm^-1^ peak intensity. **e** 1613 cm^-1^ peak intensity.

### 2.3 Environmentally-induced single-cell metabolic variations

^13^C-induced redshifts also occurred when nanoFTIR measurements were performed on cells cultured to early-exponential phase under varying concentrations of labelled glucose (n=50). Specifically, intensity of the ^13^C-amide I peak proportionally increased (in comparison to the 1655 cm^-1 12^C-amide I peak) with higher availability of labelled glucose (**Figure 4a**). When compared against the previous stable-isotope labelling experiment, where late-exponential phase cells of differing ^13^C proportions were examined (Figure 2), several spectral features differed distinctly. The ^13^C-amide I peak maximum was observed at 1627 cm^-1^ (as opposed to 1613 cm^-1^ as previously observed) and an additional shoulder was developed between 1700-1750 cm^-1^ (Figure 4a), which may be due to an increase in lipid^[42]^ or DNA concentration^[43, 44]^ in early-exponential phase cells.

**Figure 4:**
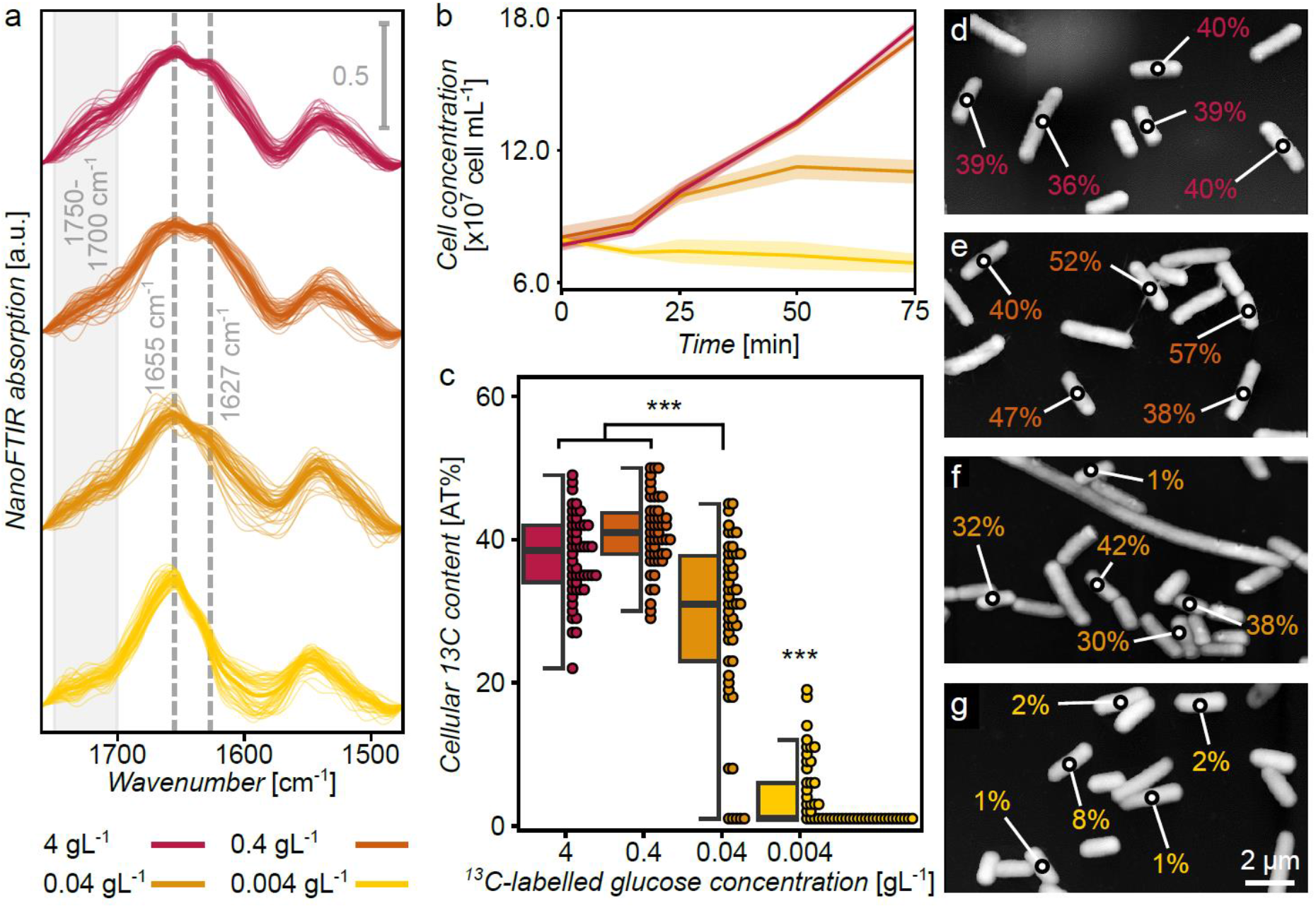
Single-cell measurements under varying concentrations of ^13^C-labelled glucose. Incremental ^13^C-labelled glucose concentration treatments are represented with a red (4 gL^-1^) to yellow (0.004 gL^-1^) color gradient. **a** NanoFTIR absorption spectra (2^nd^ harmonic) of individual single cells, with the mean spectrum of each treatment (n=50, technical replicates) shown in bold. Dashed vertical lines mark the ^12^C-(1655 cm^-1^) and ^13^C-amide I peaks (1627 cm^-1^), while the vertical gray bar marks the 1700-1750 cm^-1^ shoulder. **b** Mean *E. coli* growth curves (n=4, biological replicates), with shaded regions representing standard deviations. **c** Boxplots (displaying median, upper and lower quartiles, and the maximum and minimum excluding outliers) of quantified single-cell ^13^C content. Dots represent individual data points. Asterisks’ indicate statistical differences between treatments (p-value < 0.001 = ***), as determined by a one-way ANOVA at the 95% confidence level. **d-g** Representative AFM images of each ^13^C-labelled glucose concentration treatment. The cellular ^13^C-content, as detected by nanoFTIR is overlaid, demonstrating the variance in quantified ^13^C content of neighboring single cells. Maximum z-distances = 523, 425, 431 and 452 nm, d-g respectively.

Reducing the glucose concentration resulted in a reduction in growth rate and a corresponding decrease in the mean cellular ^13^C content. Such differences were most prominent when cells were provided with only 0.004 gL^-1^ of ^13^C-labelled glucose, with these cultures remaining in the lag phase for the entire duration of this experiment (**Figure 4b**). The lack of cellular replication is reflected in the dominant ^12^C-amide I IR absorption peak (1655 cm^-1^). This corresponded to a mean cellular ^13^C content of only 4.1%, which is >9 times less than 4 gL^-1^ glucose control samples (Q_(1, 98)_ = 30.821, p < 0.001; **Figure 4c**). However, individual cells in this glucose-limited treatment were observed with ^13^C content as high as 19% (Figure 4c). This elevated ^13^C content suggests that these few specific cells are actively replicating while the remainder of the culture is dormant. A glucose concentration of 0.04 gL^-1^ resulted in these cultures reaching carrying capacity at ∼10^8^ cell mL^-1^ (Figure 4b), which is greater than a 1-log reduction when compared to non-limited glucose cultures (Figure S3). This treatment saw the mean quantified cellular ^13^C-content reduced to 28.2% (Q_(1, 98)_ = 8.580, p < 0.001), and a dramatic increase in cell-to-cell variance (151.0; Figure 3c). In contrast, reducing the labelled glucose concentration to 0.4 gL^-1^ had a minimal to no effect. When compared to 4 gL^-1^ control samples, cellular doubling time was not influenced (Q_(1, 2)_ = 0.062, p = 0.812; Figure 4b), the spectral shape was highly similar, the mean ^13^C content quantification did not statistically differ at the 95% confidence level (F_(1, 98)_ = 3.001, p = 0.150), and the population variances were highly similar (33.4 and 24.6, respectively; Figure 4c).

These spectroscopic observations were complemented by AFM imaging (**Figure 4d-g**). Marking the ^13^C content of individual neighboring cells provides an easy visual assessment of the cell-to-cell variance between treatments and with the previous ^13^C-labelling experiment (e.g., Figure 2c). When examining morphological features, clear trends were present between cell length and glucose concentration treatments. The 4 gL^-1^ and the 0.4 gL^-1^ treatments both contained relatively equal proportions of cells typical in length (1-3 μm) and those demonstrating moderate cell elongation (3-5 μm). The proportion of cells undergoing filamentation (>5 μm in length) in these two treatments was 1.3% and 2.6%, respectively. In contrast, the 0.04 gL^-1^ and 0.004 gL^-1^ glucose concentrations were both dominated by typical length cells, with ∼77% of cells falling in this range. However, the proportion of both cell elongation and filamentation in these two samples did differ. Incubation with 0.04 gL^-1^ of glucose resulted in 18.4% elongated cells and 4.6% being >5 μm (an example of a heavily filamented cell can be seen in Figure 4f). The 0.004 gL^-1^ glucose limitation treatment saw the lowest proportion of 3-5 μm length cells of any treatment (10.7%), whereas the proportion of filamented cells exceeded this, comprising 12.4% of this population (**Figure S4**).

Furthermore, the moderate positive correlation in ^13^C amide I peak intensity with glucose concentration (Figure 4a), and the relatively high proportion of elongated cells present in the two actively replicating treatments of the glucose limitation experiment (4 gL^-1^ and 0.4 gL^-1^, Figure S4) are indicative of active cell division.^[45]^ The control sample from the glucose-limitation experiment (i.e., 4 gL^-1 13^C-labelled glucose) and the 100% ^13^C treatment of the previous experiment, only differed in incubation time, with the former being sampled at early-exponential phase and the latter in late-exponential phase. Therefore, the observed spectral differences are a result of the cellular growth phase at the time of sampling.

Correlation of several distinct spectral features (1735, 1710, 1655, 1627, 1613, 1540 and 1525 cm^-1^) with multiple environmental factors employed as treatments across the glucose limitation experiment and the previously discussed SIP labelling experiment (Figure 2), revealed three distinct subsets of data (**Figure 5**); a. both early and late-exponential phase, non-isotopically labelled cells; b. late-exponential phase, ^13^C-labelled cells; c. early-exponential phase, ^13^C-labelled cells. Furthermore, cells from the two glucose-limitation treatments that were seen to affect growth (0.04 and 0.004 gL^-1^) each showed spectral similarities with both the ^12^C cluster and the other early-exponential phase samples (Figure 5, orange and yellow dashed circles, respectively).

**Figure 5:**
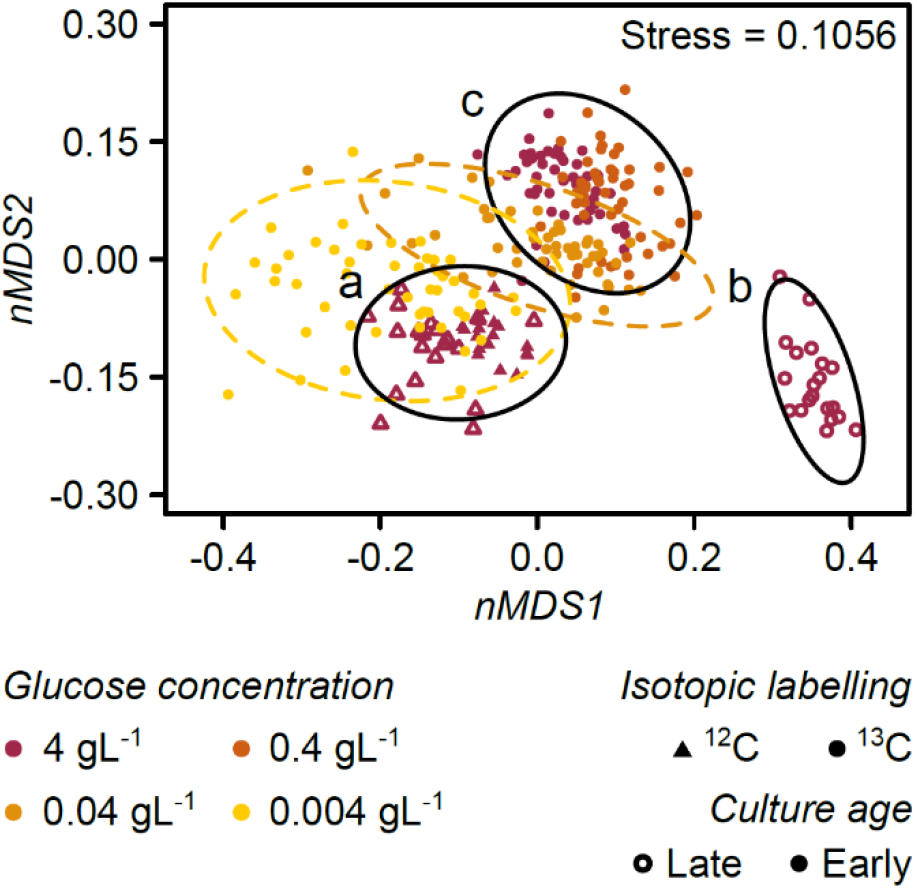
Non-metric multidimensional scaling (nMDS) plot based on normalized nanoFTIR absorption intensities at seven key spectral positions, measured on single *E. coli* cells cultured under differing ^13^C-labelled glucose treatments. Incremental ^13^C-labelled glucose concentration treatments are represented with a red (4 gL^-1^) to yellow (0.004 gL^-1^) color gradient. ^13^C labelling is represented by symbols (circles = 100%, triangle = 0%), and cellular growth phase at the time of harvest is represented by the symbol fill (solid = early-exponential, outline = late-exponential). Ellipses represent 95% groupings, with black defining distinct datasets (a-c), and colored-dashed ellipses defining datasets that span multiple groupings.

Further investigation of growth-phase dependent spectral features was performed, comparing three different examples of cultures, each composed of 50% ^13^C-labelled proteins (**Figure 6a**). These three unique conditions each resulted in different subcellular isotope distributions, that subsequently effected the extent of ^13^C-^13^C mode coupling and its influence on the resulting amide I peak shape (**Figure 6b**). Specifically, the theoretical 50% spectrum (derived from a linear combination of 0% and 100% ^13^C-labelled, late-exponential growth phase cells) resulted in equal contributions from non-labelled and completely stable-isotope labelled proteins. This heterogenous protein composition provides an example of maximal ^13^C-^13^C-mode coupling, which results in the ^12^C-amide I peak dominating the ^12^C:^13^C peak ratio (Figure 6b, gray). This is contrasted by the experimentally derived, late-exponential growth phase cells, which contained a relatively homogenous isotope distribution, brought about due to prolonged exposure to 50% ^13^C-labelled media. The decreased ^13^C -^13^C interaction in this sample resulted in an approximately equal ^12^C:^13^C amide I peak ratio (Figure 6b, light purple). Lastly, in addition to the unique shoulder present at ∼1730 cm^-1^ (as described in Figure 4a), early-exponential phase samples (composed of cells supplied with 100% ^13^C-labelled glucose and harvested after approximately one doubling time) had an amide I shape with similarities to both other examples (Figure 6b, dark purple). This suggests that these samples are comprised from a mixed isotope distribution model, showing reduced but still present ^13^C-^13^C mode coupling.

**Figure 6:**
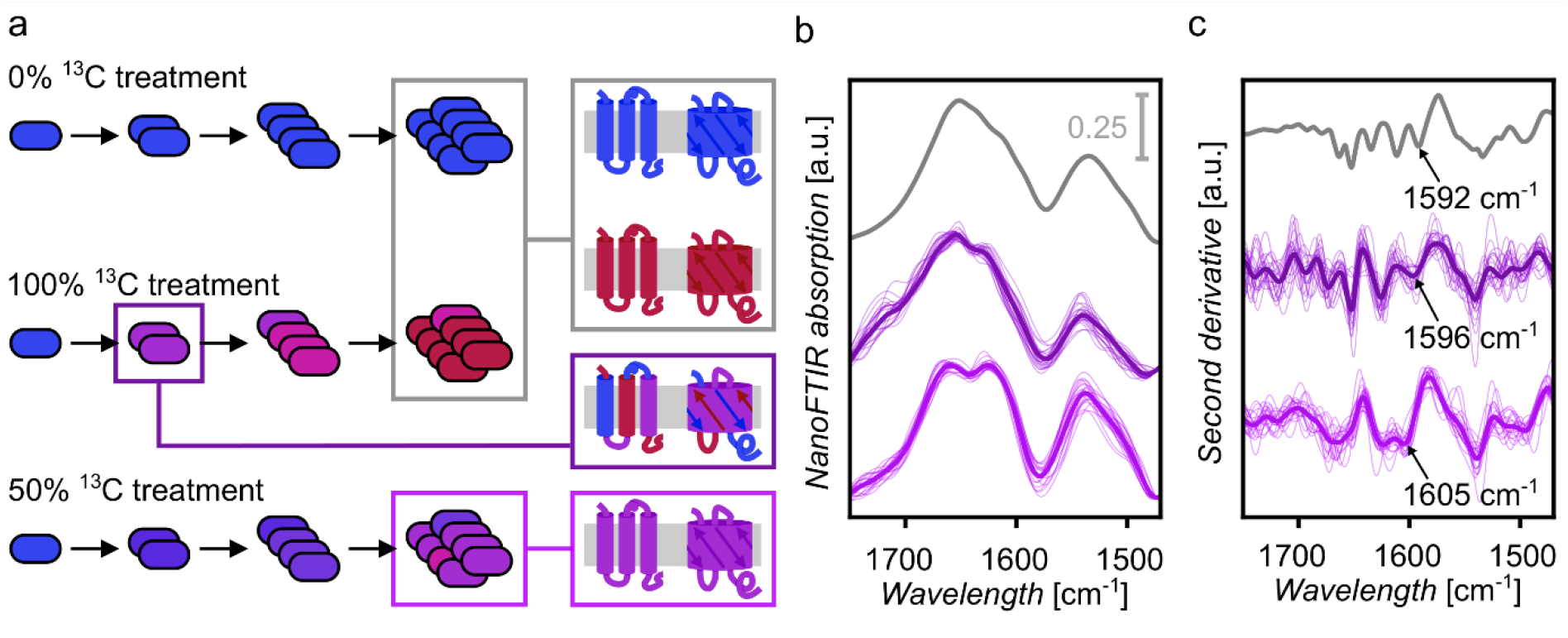
Assessment of sub-cellular isotopic conformations. **a** An illustrative diagram of cellular replication under three different culture conditions, and the resulting theoretical isotopic composition of protein secondary structures. Proportions of ^13^C are represented with a blue (0%) to red (100%) color gradient. **b** NanoFTIR absorption spectra (2^nd^ harmonic) and **c** second derivative plots of the amide region of three variants of 50% ^13^C-labelled samples; a theoretical spectrum derived from a linear combination of mean 0% and 100% late-exponential spectra (gray), the early-exponential experimental samples (dark purple), and the late-exponential experimental samples (light purple), with the means of the latter two treatments (n=20, technical replicates) shown in bold.

Although the amide I peak is complex in nature, being derived from multiple proteins, each in various orientations and conformations, the spectral contributions of different secondary structures (α-helices, β-sheets and turns) and their respective ^13^C-induced redshifts, are well studied.^[41, 46-48]^ Second derivative analysis, a common tool to detect hidden peaks within broad spectral features,^[49-51]^ was therefore utilized to differentiate amide I secondary structure compositions (**Figure 6c**). Second derivate analysis of unlabeled and 100% ^13^C-labelled cells showed distinct and distinguishable components of the amide I peak. As expected, α-helices (^12^C = 1652 cm^-1, 13^C = 1613 cm^-1^) and β-sheets (^12^C = 1664 cm^-1^ & 1634cm^-1, 13^C = 1592 cm^-1^) were the major contributors to the overall amide I peak shapes (**Figure S5**).^[52]^ In contrast, several compositional differences were present in the second derivative plots of the three different 50% labelled samples (**Figure 6c**). The second derivative plot of the theoretically calculated 50% sample was comprised of several clear and distinct features that corresponded to the above assignments (Figure 6c, gray). The remaining samples, with increased isotope homogeneity (and hence decreased ^13^C-^13^C mode coupling) saw high levels of cell-to-cell variation. Many previously resolved features are more ambiguous in the mean second derivative plot of the late-exponential phase sample (Figure 6c, light purple). Despite this, several assigned peaks are distinct across all three samples and remain unaffected by neighboring peaks. Specifically, the ^13^C-β-sheet peak shifts from 1592 cm^-1^ in the theoretically derived sample, to 1596 cm^-1^ in the mean early-exponential sample, to 1605 cm^-1^ in the mean late-exponential phase sample. This increase in spectral position of the ^13^C-β-sheet is due to the decrease in ^13^C-^13^C mode coupling of the samples examined, and is similar to previously observed correlations in wavenumber and mode coupling.^[47]^ A similar shift was also in the ^13^C-α-helix peak (1613-1626 cm^-1^), however this assignment is less prominent due to some spectral overlap with the ^12^C-β-sheet peak. Additionally, second derivative analysis also revealed that the amide I shoulder unique to early-exponential phase cells was comprised of two peaks positioned at 1718 cm^-1^ and 1740 cm^-1^ (Figure 6c, dark purple). It is possible that these peaks are arising due to specific label symmetries (i.e., the distance-relationship of isotopes both within and between biomolecules) within the proteins^[46]^ or lipid bilayers.

## 3. Discussion

This study introduces and demonstrates SIP-nanoFTIR as a robust and powerful new tool capable of simultaneous quantification of single-cell metabolism and acquisition of high-resolution morphological information. When used to examine ^13^C-labelled *E. coli* cells the observed amide I redshift was used to determine the carbon isotopic enrichment of newly synthesized proteins and hence single-cell translational activity. In general, IR-absorption recordings are quick, cost effective and easy to perform, however, averaged measurements derived from a large number of cells inherently results in a loss of fine-scale information. Near-field IR nanoscopy overcomes this limitation, and SIP-nanoFTIR measurements yielded comparable spectra and similar mean cellular ^13^C content to well-established IR measurements of bulk cell-numbers (such as IR-reflectance microscopy) and to Raman-based studies.^[34, 35]^

SIP-nanoFTIR is highly suited to investigations of intercellular heterogeneity, with the results of this work corresponding well with other recent stable-isotope labelling studies, including observations of similar levels of cell-to-cell heterogeneity using a combination of mid-IR photothermal microscopy and fluorescence *in-situ* hybridization (FISH).^[1]^ Glucose limitation emphasized intercellular competition, which induced population-wide changes in cellular morphology and was a driver of cellular metabolic heterogeneity. Some outlier cells had dramatically elevated ^13^C levels while others acquired little to no additional carbon. These results are in accord with previous experimental observations using single cell SIP-nanoSIMS that also identified nutrient limitation as a driver of heterogeneous substrate assimilation.^[53]^ Variation in glucose scavenging rates at the single-cell level means that individual outlier cells with elevated metabolic rates are significantly outcompeting the majority of the population, and as such may play a disproportionately large role in shaping the structure and function of growing microbial communities. SIP-nanoFTIR therefore provides a comprehensive means to link single-cell metabolic processes to population-wide changes originating in response to environmental fluctuations. In a similar fashion, other isotope-labelling based analytical techniques have also provided insights into the ecological function of the individual cells that comprise complex microbial communities.^[1, 35, 54]^ Additionally, quantification of cellular heterogeneity using SIP-nanoFTIR has applications in biopharmaceutical and medicinal fields, complementing studies such as the characterization of high-value eukaryotic proteins,^[55]^ the determination of patient-specific drug effectivity,^[56]^ *in-situ* detection of antibiotic-resistant bacteria or biofilms,^[57, 58]^ and the metabolic tracing of cancer cells^[59, 60]^ or the human microbiota.^[61, 62]^

Isotopically induced changes in the amide I peak shape arose both in response to environmental factors and were dependent on the cellular growth phase of harvested cultures. A higher number of cellular replication cycles resulting in a more homogenous isotope distribution within newly synthesized proteins, which decreased ^13^C-^13^C vibrational mode coupling changes. Consequently, spectral changes were observed in amide I secondary structures, most prominently due to isotopic-label symmetries within antiparallel beta-sheets.^[46]^ Analysis of subcellular isotopic heterogeneity provides evidence that proteins synthesized during early-exponential growth phase consist mainly (but not exclusively) of freshly metabolized carbon, rather than carbon recycled from the parent cell. Identification of such ^13^C-^13^C mode coupling induced features means SIP-nanoFTIR, in addition to single cell metabolic quantification, has the capability to spectroscopically differentiate specific cellular growth phases. SIP-nanoFTIR improves on FTIR spectroscopy, an already valuable tool for the monitoring of microbial growth phases, alleviating several of the constraints identified.^[63]^

Future SIP-nanoFTIR investigations should consider other stable isotopes (for instance, ^2^H or ^15^N)^[34, 38, 40, 58, 64, 65]^ and explore cellular metabolism under differing experimentally manipulated environmental stressors.^[63]^ The combination of spectroscopic and phenotypic information derived from such studies could provide links between genetic adaptations, induced changes in morphology and molecular composition, and cellular function. Observations of subcellular biomolecule localization and transport via stable-isotope labelling would be most effective in more complex eukaryotic organisms. Both nanoFTIR and sSNOM have been used to image eukaryotic samples prepared via thin sectioning.^[25]^

Combining this method with stable-isotope labelling would enable the acquisition of 3D-topographic and spectroscopic information, dictated by localized, internal cellular isotope distributions and the cell’s metabolic contributions to specific membrane-bound organelles. Additionally, SIP-nanoFTIR can be effectively combined with several complementary or correlative single-cell techniques, including nanoSIMS,^[10]^ high resolution Raman spectroscopy,^[66]^ and FISH.^[1]^ Many of these techniques have already been used together,^[40]^ even examining the same individual cell.^[54]^ Pairing these correlative techniques with SIP-nanoFTIR could greatly enhance our understanding of complex cellular processes and dynamics, and may be particularly relevant for analyzing complex samples such as environmental communities or samples from multi-cellular organisms. The data redundancy yielded through correlative techniques can be used as a means of validation, enhancing reliability and further strengthening the experimental outputs of all techniques involved. Finally, an ongoing challenge for all single-cell techniques is data acquisition time and consequently, sample size. However, recent advancements in automated and AI-supplemented sampling offer promising solutions,^[67, 68]^ making high-throughput single-cell measurements a feasible and realistic goal.

This work effectively demonstrates SIP-nanoFTIR as a highly advantageous near-field technique that effectively revealed small-scale, cell-to-cell variations that traditional bulk cell analysis techniques fail to capture. SIP-nanoFTIR offers the advantage of not requiring a large number of cells, making it particularly suitable for examining environmental samples containing naturally low cell numbers or studying microorganisms with slow metabolic activity in their native habitats. The simultaneous acquisition of high-resolution topographic information and quantification of single-cell biosynthesis further enhances our understanding of environmental responses. Insights into the dynamics of single-cell microbial competition and metabolism can be used in identifying outlier organisms that may exert a disproportionate influence on shaping population dynamics. Given the wide range of potential applications, SIP-nanoFTIR is expected to provide valuable insights in various fields, including microbial ecology, cell biology, biophysics, and specific medical science applications. Its ability to explore cellular heterogeneity and dynamics with high sensitivity and resolution promises to deepen our understanding of fundamental biological processes and pave the way for novel discoveries in numerous scientific disciplines.

## 4. Methods

### Culture conditions

To eliminate any trace carbon, prior to autoclave sterilization, all glassware was heated to 500 °C for 3 h. *Escherichia coli* K12 (DSM 498) were aerobically cultured at 37 °C in M9 minimal medium composed of 10.5 gL^-1^ M9 Broth (Merek, Germany), 1.5 mM MgSO_4_, and 0.1 mM CaCl_2_ supplemented with a single carbon source; 4.0 gL^-1^ of D-glucose provided in specific isotopic ratios (^12^C, Merek, Germany; ^13^C, Cambridge Isotope Laboratories, USA; Table S1). 10^9^ cells were harvested at late-exponential growth phase (Figure S3), fixed in 2.5% (v/v) glutaraldehyde (Grade I, Merek, Germany) at room temperature for two hours, washed and resuspended in milliQ water.

Additional samples of *E. coli* were cultured under limited carbon conditions; M9 minimal media (as described above) was supplemented with ^13^C-labled glucose, provided at concentrations of 4.0, 0.4, 0.04 and 0.004 gL^-1^. Due to the distinct observational goals of this glucose-concentration experiment, unlike the isotope ratio experiment, cells were harvest during the early-exponential phase (Figure 4b). Cells were then fixed, washed and resuspended in the same manner described above.

### Single-cell spectroscopy and imaging

2 μL cellular samples were dried onto a template stripped gold substrate, and for each varied isotope proportion treatment, single-cell nanoFTIR spectra were acquired from the center of 20 randomly selected individual cells, using the spectroscopy module of a neaSNOM Nearfield Microscope (Neaspec, Germany). The nanoFTIR spectrometer was operated in tapping mode using coated AFM probe tips (Arrow NCPt, NanoWorld, Switzerland) with a resonance frequency of ≃285 kHz. A 2050-1010 cm^-1^ broadband laser source was used, and spectra were recorded with a resolution of 12 cm^-1^. Each spectrum was derived from 100 scans with a 6 ms integration time, and was compared against a spectrum of the blank gold substrate. For samples cultured under glucose-limited conditions, spectra were acquired in the same manner from 50 randomly selected cells per treatment. NanoFTIR spectra were linearly baseline corrected, and the FTIR-absorption was calculated from combined phase and amplitude of the near-field signal.^[69]^ When required, spectra were smoothed using a Savitzky-Golay filter at window size 11, polynomial order 2. AFM images were acquired using the nanoFTIR spectrometer set up as described above, operated in tapping mode. Visualization and analysis were performed with Gwyddion (v.2.62, Czech Metrology Institute, Czech Republic).

NanoFTIR hyperspectral mapping was performed on a single, dried *E. coli* cell originating from a culture incubated with 4 gL^-1^ of 50% ^13^C-labelled glucose. A total of 312 spectra were acquired over the surface of the sample, with a spatial resolution of 100 nm/spectrum. To account for sample drift, a single unaveraged spectrum was acquired at each sample position, then the samples was reoriented to the exact same starting position (using AFM imaging). A total of 50 complete scans were performed, each referenced against a new background measurement. Manual spectral averaging was performed at each of the 312 positions, providing adequate signal to noise. Spatial masking was performed based on a threshold value of 10.5 V of the unreferenced amplitude signal of the scattered laser light.

### IR-reflectance spectroscopy

While SIP-nanoFTIR and IR-reflectance measurements could not be performed on the exact same cellular preparations (due to the higher cellular concentration required for IR-reflectance microscopy resulting in overcrowding of SIP-nanoFTIR samples), great care was taken to ensure the only difference between samples was the cellular concentration; each sample was prepared from the same clonal population of fixed *E. coli* cells and deposited onto the same template-strip gold surface. IR-reflectance microscopy was performed in triplicate for each treatment, with spectra acquired from an average of 65 equally distributed points (Figure S2), using an IR-reflectance microscope (Hyperion 2000, IR source proved by a Vertex 80v, Bruker, Germany) fitted with a 15x Cassegrain objective (NA=0.4, Newport, Germany), performing 500 scans at each point, from 650-3950 cm^-1^ at 4 cm^-1^ resolution.

### IRMS

For IRMS analysis samples were prepared in triplicate by depositing fixed cells onto 25 mm diameter glass fiber filters (0.7 μm pore size, Whatman, UK; trace carbon sterilized at 500 °C, 3 h). Following cellular deposition, filters were washed with an excess of milliQ water, air dried, then exposed to 20% HCl vapor for >12 h. Five 5 mm diameter subsamples were punched from each filter and wrapped in aluminum capsules (3.5mm × 5mm, HEKAtech, Germany). Subsamples were combusted to CO_2_ gas using an elemental analyzer (Flash 2000 CHNS/O, Thermo Fisher Scientific, Germany), performed with a 10 mL O_2_ pulse added to a 1050 °C combustion chamber filled with wolfram oxide and silver-cobalt oxide. Using helium carrier gas (gas flow = 0.08 Lmin^-1^), combustion product gases were swept through a 650 °C reduction furnace filled with copper. Remaining water was trapped with phosphorus pentoxide. Gases were isothermally separated at 70 °C on a gas chromatography column (HE26070500, HEKAtech, Germany) and transferred to the IRMS via an open split system (ConFloIV, Thermo Fisher Scientific, Germany). The carbon isotopic composition (expressed as AT%) was normalized with a two-point calibration against ^12^C D-glucose (1.09 AT%) and ^13^C D-glucose (99 AT%) standards, yielding an analytical precision of ±0.7 AT%.

### Cellular ^13^C quantification

In order to utilize observed spectral differences to calculate single-cell ^13^C content, the isotopically-induced amide I redshift was used to formulate a quantification algorithm. By focusing solely on the ratio of peak intensities at two positions, the influence of other, less relevant spectral features (such as locally elevated lipid concentrations or near-field enhancement of membrane protein secondary structures) was minimized. Two calibration steps were used to translate spectral amide I information into ^13^C AT%. Firstly, IR-reflectance spectra of organisms provided with 10% incremental ^13^C portions (from 0% to 100%) were measured. This far-field spectroscopy provided the mean spectral information from a large number of organisms (in the order of 10^7^ cells), while near-field induced peak shifts with respect to the far-field data were accounted for by individually assigning ^12^C and ^13^C amide I peak positions for each method. The amide I redshift can be enhanced by both the number and the symmetry of isotopic labels,^[70-72]^ and as a result proportional ^13^C incorporation does not linearly correlate with shifting peak intensity. The resulting peak intensity ratios at 1655 cm^-1^ (^12^C amide I) and 1613 cm^-1^ (^13^C amide I) declined from ∼3:1 (0% treatment) to ∼1:2 (100% treatment) in a 3^rd^ order polynomial curve. Secondly, IRMS measurements of cells cultured under 25% ^13^C increments (from 0% to 100%) were used to compare the proportion of ^13^C provided with the mean cellular ^13^C AT%. This curve was fitted using a second order polynomial. Using these two fit-functions allows for an automated translation from each nanoFTIR spectrum to the single-cell metabolized ^13^C (Figure S1). To account for spectral differences in early-exponential growth cells, a linear combination of averaged 0% and 100% nanoFTIR spectra was used (simulating comparable ^13^C-^13^C vibrational mode coupling). Peak ratios at 1655 cm^-1^ (^12^C amide I) and 1613 cm^-1^ (^13^C amide I) declined from ∼3:1 (0% treatment) to ∼1:2 (100% treatment), in a 3^rd^ order polynomial curve.

### Data analysis

The mean quantified ^13^C content from cells cultured under differing ^13^C fractions of supplied ^13^C-labelled glucose was compared using one-way analyses of variance (ANOVA) with Tukey’s post hoc tests. This was performed using both nanoFTIR measurements (Table S2) and IR-reflectance microscopy (Table S3) as categorical variables. One-way analyses of variance (ANOVA) with Tukey’s post-hoc tests were also used to compare the mean quantified ^13^C content of each glucose-limited treatment, as well as both the cellular doubling time and cell concentration at the conclusion of the incubation period. To construct the nMDS plot, a Bray-Curtis dissimilarity matrix was calculated based on the normalized peak intensity at 1735, 1710, 1655, 1627, 1613, 1540 and 1525 cm^-1^, with three discrete variables considered; ^13^C labelling, glucose concentration and the growth phase at the time of cellular harvesting. Specific datasets were grouped based on 95% confidence intervals. Visualization and statistical analyses were performed with RStudio (v.2023.12.0+369, Postit Software, USA).

## Acknowledgements

D.J.B. and J.D. contributed equally to this work. This work gratefully acknowledges financial support from the Bundesministerium für Wirtschaft und Energie (Projektträger Deutsches Zentrum für Luftund Raumfahrt) grants 50WB1623 and 50WB2023 (A.E., D.J.B.), the Volkswagen Foundation and its Freigeist Program (A.E.), and the German Research Foundation DFG project number 462858357 (A.P.). We further acknowledge support from the SupraFAB research building realized with funds from the federal government and the city of Berlin, and the analytical facilities of the Centre for Chemical Microscopy (ProVIS) at UFZ Leipzig, which is supported by the European Regional Development Funds (EFRE-Europe funds Saxony) and the Helmholtz Association, and the Laboratories for Stable Isotopes (LSI) at UFZ Leipzig for the use of their instrumental resources.

