## Supporting Information for "Stable Isotope Probing-nanoFTIR for Quantitation of Cellular Metabolism and Observation of Growth-dependent Spectral Features"

### These authors contributed equally

* Corresponding authors

**Table S1: IRMS measurements of cellular standards**.

Samples cultured under varying proportions of supplied $^{13}C$-labelled glucose.

| Supplied $^{13}C$ fraction [%] | Mean cellular $^{13}C$ fraction [AT%] | Standard deviation [AT%] |
|---|---|---|
| 0 | 1.1 | 0 |
| 25 | 25.29 | 0.02 |
| 50 | 46.17 | 0.35 |
| 75 | 64.62 | 0.39 |
| 100 | 79.61 | 0.23 |

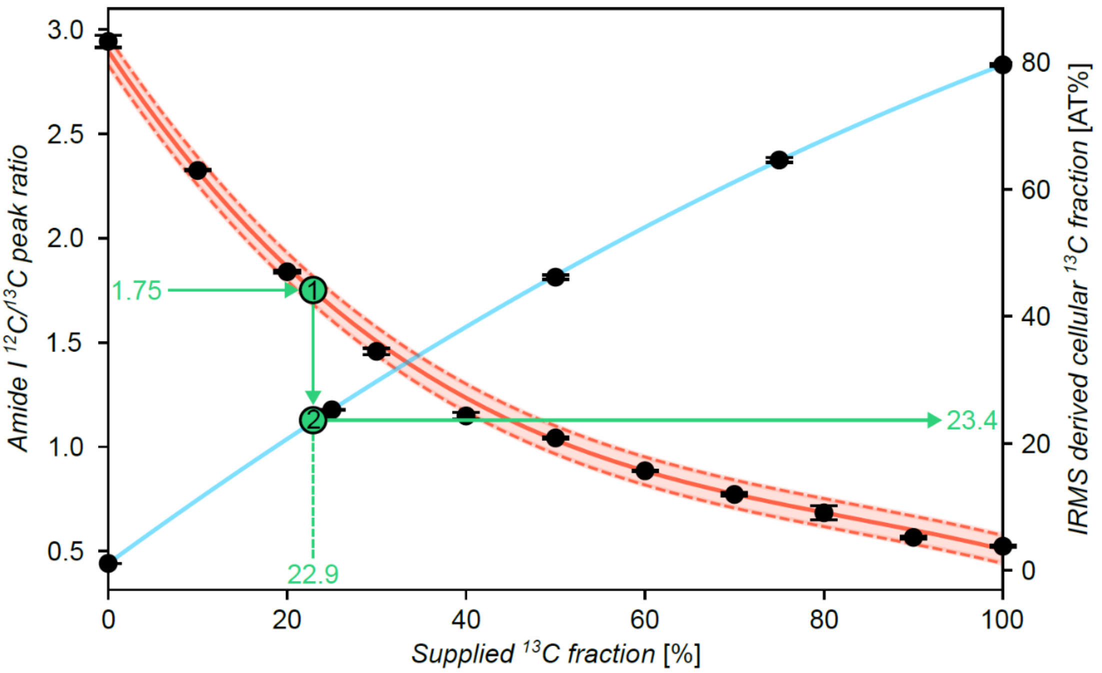


**Figure S1: NanoFTIR-based quantification of cellular $^{13}C$ content, following IRMS standardization.**

Quantification was performed by applying a two-step algorithm:

1. $\frac{12C\ peak\ height}{13C\ peak\ height} = f_{1/2}(treatment)\ \rightarrow treatment = f_{1/2}^{-1}$

2. $AT\% = f_3(treatment)$

Specific polynomial equations were applied to each experimental group as follows:

1. Third order polynomial applied to late-exponential cells

$$f_1(x) = -2.657 \cdot 10^{-6} \cdot x^3 + 6.665 \cdot 10^{-15} \cdot x^2 - 0.064 \cdot x + 2.896$$

2. Third order polynomial applied to early-exponential cells

$$f_2(x) = -2.531 \cdot 10^{-6} \cdot x^3 + 6.876 \cdot 10^{-4} \cdot x^2 \pm 7.376 \cdot 10^{-2} \cdot x + 3.524$$

3. Second order polynomial applied to IRMS measurements

$$f_3(x) = -0.002 \cdot x^2 + 1.028 \cdot x + 1.100$$

**Table S2: Statistical comparison of the mean cellular $^{13}$C content, measured by nanoFTIR.**

Cells cultured under five different fractions of $^{13}$C-labelled glucose (0, 25, 50, 75, 100%) were compared at the 95% confidence level, using a one-way ANOVA (top) and Tukey's post-hoc test (bottom).

| *ANOVA Score* | *Sum of Squares* | *df* | *Mean Squares* | *F* | *p-value* |
|---|---|---|---|---|---|
| *Between Groups* | 4 | 76471.3 | 19117.83 | 2393.823 | <0.00001 |
| *Within Groups* | 95 | 758.7 | 7.9863 | | |
| *Total* | 99 | 77230 | 780.101 | | |

| *Tukey's HSD* | *Mean Difference* | *Standard Error* | *p-value* |
|---|---|---|---|
| *0% v 25%* | 27.45 | 0.6319 | <0.00001 |
| *25% v 50%* | 17.3 | 0.6319 | <0.00001 |
| *50% v 75%* | 20.95 | 0.6319 | <0.00001 |
| *75% v 100%* | 12.15 | 0.6319 | <0.00001 |

**Table S3: Statistical comparison of the mean cellular $^{13}$C content, measured by IR reflectance microscopy.**

Cells cultured under five different fractions of $^{13}$C-labelled glucose (0, 25, 50, 75, 100%) were compared at the 95% confidence level, using a one-way ANOVA (top) and Tukey's post-hoc test (bottom).

| *ANOVA Score* | *Sum of Squares* | *df* | *Mean Squares* | *F* | *p-value* |
|---|---|---|---|---|---|
| *Between Groups* | 4 | 11215.334 | 2803.833 | 539.1987 | <0.00001 |
| *Within Groups* | 10 | 52 | 5.2 | | |
| *Total* | 14 | 11267.334 | 804.8095 | | |

| *Tukey's HSD* | *Mean Difference* | *Standard Error* | *p-value* |
|---|---|---|---|
| *0% v 25%* | 29.6667 | 1.3166 | <0.00001 |
| *25% v 50%* | 17 | 1.3166 | 0.00002 |
| *50% v 75%* | 20.6667 | 1.3166 | <0.00001 |
| *75% v 100%* | 9 | 1.3166 | 0.00482 |

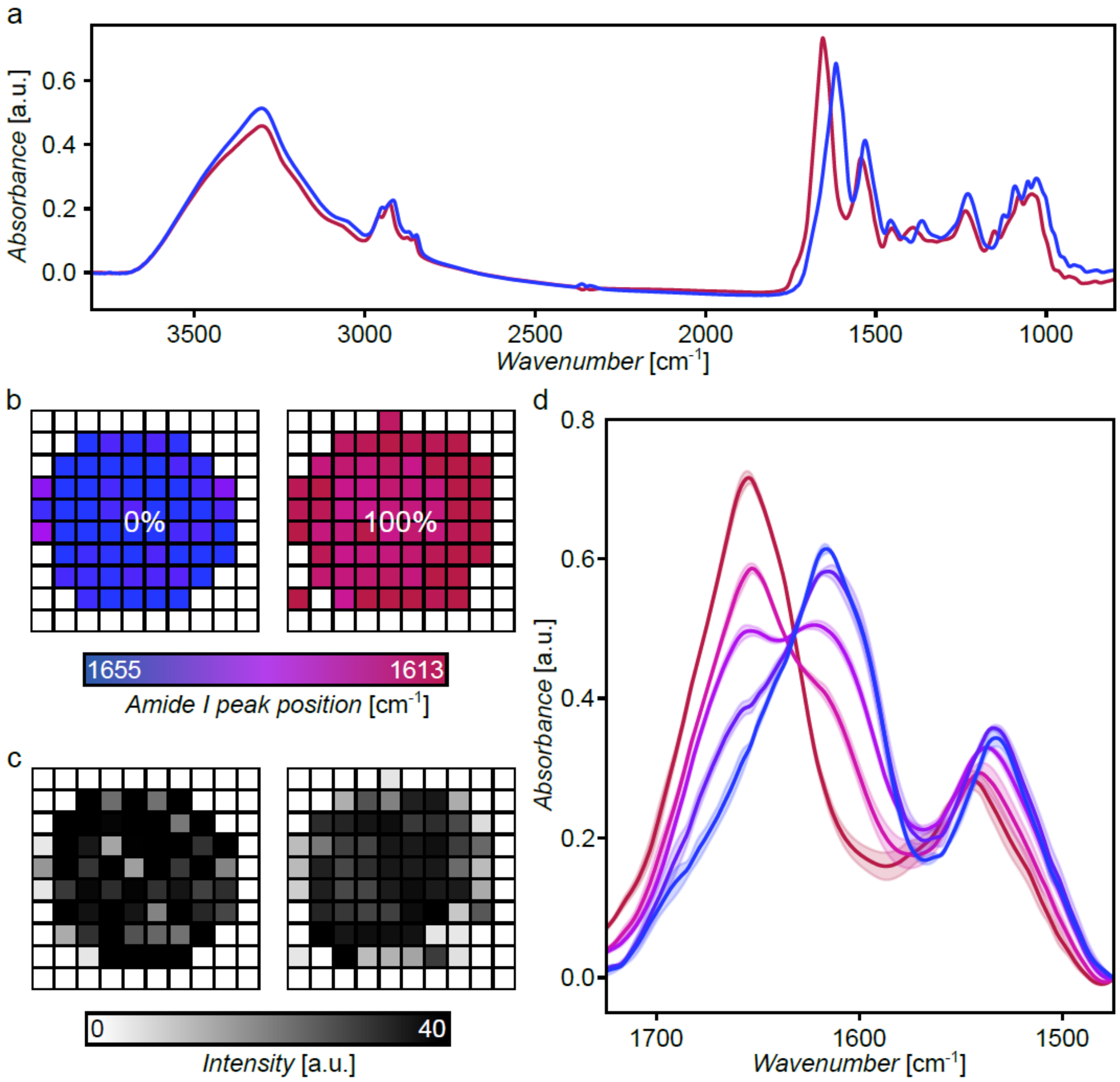


**Figure S2: IR-reflectance microscopy measurements.**

25% incremental $^{13}$C-labelled glucose treatments are represented with a blue (0%) to red (100%) color gradient. **a** Complete IR absorbance spectra of 0% and 100% labelled cells. **b** Amide I peak position acquired from approximately 65 equally distributed points across the surface of a 2 μL droplet on a template stripped gold surface, from representative samples of the 0% and 100% $^{13}$C-labelled treatments. Each pixel represents ~0.0625 mm$^2$. **c** Amide I peak intensity corresponding to the position examined above. **d** Mean IR absorbance spectra of the amide region of five different mean $^{13}$C-labelled samples (n=3, technical replicates), derived from average of intensity-weighted spectra across the droplet surface. Shaded regions represent standard deviations.

32

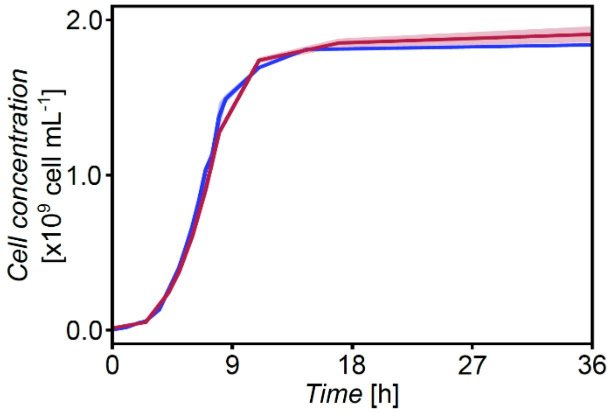


**Figure S3: Mean growth curves of stable-isotope labelled *E. coli* (n=2, biological replicates).**

Cells were cultured under growth media supplemented with 0% (blue) and 100% (red) $^{13}$C-labelled glucose. Shaded regions represent standard deviations.

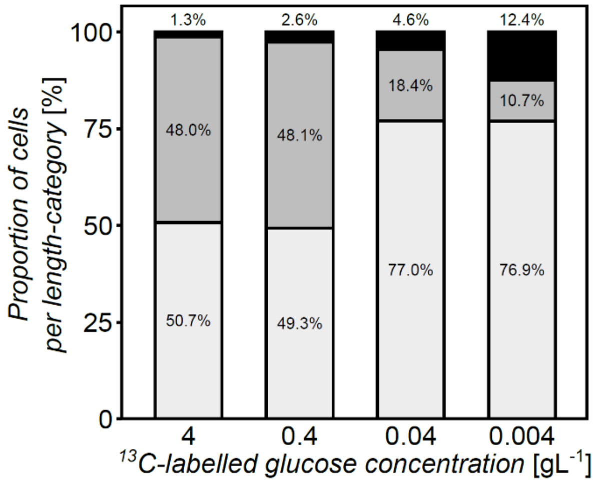


**Figure S1: Proportions of *E. coli* cell lengths present in samples with varying $^{13}$C-labelled glucose concentrations (n>225, technical replicates).**

Cell length categories were divided into three categories: <3 µm (white), 3-5 µm (gray) and >5 µm (black).

33

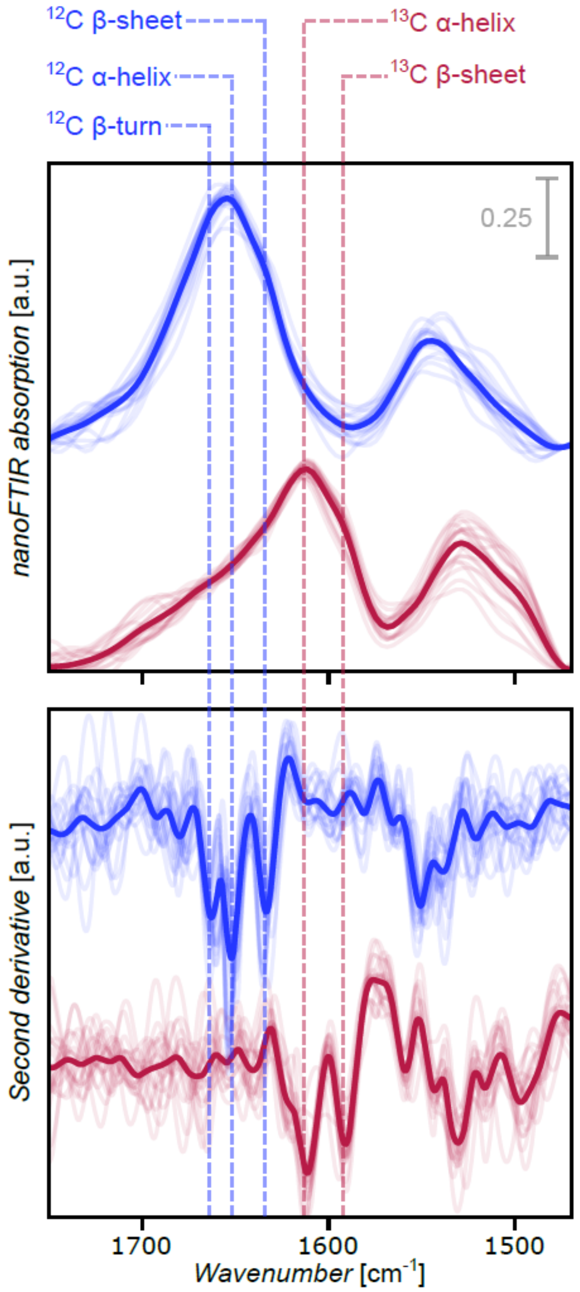


**Figure S2: Amide spectra (top) and second derivative plots (bottom) of individual unlabelled (blue) and 100% $^{13}$C-labelled (red) *E. coli* cells.**

The mean nanoFTIR absorption spectrum (2$^{nd}$ harmonic) and second derivative plot of each treatment (n=20, technical replicates) are shown in bold. Cells are harvested and fixed at late-exponential growth phase. Dashed vertical lines mark the peak assignments of five specific secondary protein structures.
